# Biocompatible Membrane Vesicles from *Lactobacillus acidophilus* MTCC 10307 Exhibit Potent Anti-Inflammatory Activity

**DOI:** 10.64898/2026.04.01.715785

**Authors:** Venkatramanan Mahendrarajan, Nalini Easwaran

## Abstract

Inflammation is a fundamental immune response but, when dysregulated, contributes to the pathogenesis of numerous inflammatory disorders. Although there are several conventional anti-inflammatory drugs which are effective, their long term use is often associated with adverse side effects, which highlights the need for safer alternative therapeutic drugs. Probiotic derived membrane vesicles (MVs) have recently emerged as biologically active nanostructures capable of modulating host immune responses. In the present study, MVs isolated from *Lactobacillus acidophilus* MTCC 10307 were evaluated for their anti-inflammatory efficacy and safety profile using *in vitro* and *in vivo* models. In RAW 264.7 macrophages, *L. acidophilus* MVs significantly attenuated lipopolysaccharide induced expression of the pro-inflammatory mediators *Il-1β*, *Il-6*, and *iNOS*, accompanied by reduced nitric oxide and reactive oxygen species production which was abolished in the proteinase K treated MVs. The protein levels of NFκB and IL1β were also reduced in the treatment groups. Repeated dose oral toxicity studies revealed no adverse effects, as evidenced by body weight and histopathological evaluation of major organs. The anti-inflammatory properties of *L. acidophilus* MVs were further validated in an *in vivo* hind paw edema model, which shows inflammation resolution demonstrated by molecular and histological analysis. Proteomic analysis using LC-MS/MS identified the presence of surface-layer protein A (SlpA) which is a potential bioactive component which might contribute to the observed immunomodulatory effects. Collectively, these findings demonstrate that *L. acidophilus* MVs exert potent anti-inflammatory activity while maintaining an excellent safety profile using integrated *in vitro* and *in vivo* models.

## Introduction

Inflammation is a crucial biological response that promotes host defense against infection and tissue injury (1). However, uncontrolled or unresolved inflammation may escalate into a wide range of pathological conditions, including chronic inflammatory systemic diseases, metabolic disorders, and inflammatory conditions of the oral cavity (2). At the cellular level, macrophages are primary mediators of inflammatory responses and serve as the first line of defense against microbial invasion. Upon stimulation by bacterial components such as lipopolysaccharide (LPS), macrophages initiate intracellular signalling cascades that result in the production of pro inflammatory cytokines, reactive oxygen species (ROS), and nitric oxide (NO) (3).

Among the key signalling pathways involved in inflammation, nuclear factor-κB (NF-κB) plays a central role in regulating immune and inflammatory responses (4). Activation of NF-κB leads to transcriptional upregulation of several pro-inflammatory genes, including *Il-1β*, *Il-6*, and *iNOS*. Persistent activation of NF-κB signalling has been strongly associated with chronic inflammation and tissue damage, making it an important therapeutic target for anti-inflammatory interventions (5).

Although non-steroidal anti-inflammatory drugs (NSAIDs) such as diclofenac are widely used to manage inflammation, long term administration is often associated with adverse gastrointestinal, renal, and cardiovascular effects (6). Consequently, there is developing research interest in identifying safer, biologically derived alternatives that can effectively modulate inflammatory pathways without causing systemic toxicity. Probiotic species have gained considerable spotlight for their immunomodulatory properties (7). Beyond live bacterial cells, probiotic derived components such as membrane vesicles (MVs) are now recognized as key mediators of host microbe communication (8). MVs are nano-sized lipid bilayered structures released by bacteria that encapsulate proteins, lipids, and signalling molecules. Importantly, bacterial MVs can interact directly with host immune cells and influence inflammatory signalling while avoiding the safety concerns associated with administering live bacteria.

Multiple *Lactobacillus* species have been extensively identified to exert anti-inflammatory effects through modulation of host immune signalling pathways. *Lactobacillus rhamnosus* GG has been shown to suppress LPS-induced production of pro-inflammatory cytokines such as IL-1β, IL-6, and TNF-α by inhibiting NF-κB activation in macrophages and intestinal epithelial cells (9). Similarly, *Lactobacillus plantarum* strains have been shown to reduce inflammatory responses by downregulating NF-κB and MAPK associated signalling, which decreased the expression of iNOS and COX-2 and nitric oxide production (10). Anti-inflammatory properties have also been reported for *Lactobacillus casei*, which attenuates cytokine secretion and oxidative stress in immune cells by modulating Toll-like receptor associated molecular pathways (11). Remarkably, *Lactobacillus acidophilus* and its components, including surface-layer proteins, have been shown to suppress LPS induced inflammatory mediator expression through disturbance of NF-κB and MAPK signalling cascades (12). Altogether, the listed studies indicate that suppression of NF-κB driven cytokine production, reduction of oxidative and nitrosative stress, and modulation of innate immune signalling are innate anti-inflammatory mechanisms of action across multiple *Lactobacillus* species, supporting their therapeutic capabilities in inflammatory disorders.

*L. acidophilus* is a well-established probiotic bacterium with documented anti-inflammatory properties. Among its bioactive components, surface layer protein A (SlpA) has been reported to suppress LPS-induced inflammatory responses in macrophages through modulation of TLR4 dependent MAPK and NF-κB pathways, as well as NOD2/NLRP3-associated signalling (12). However, despite increasing interest in probiotic derived vesicles, the anti-inflammatory efficacy and systemic safety of *L. acidophilus* MVs remain insufficiently explored.

Therefore, the present study aimed to comprehensively evaluate the anti-inflammatory potential of *L. acidophilus* membrane vesicles using *in vitro* macrophage models and *in vivo* rat models, while simultaneously establishing their safety profile through repeated-dose oral toxicity assessment.

## Materials and Methods

### Isolation and Characterization of *L. acidophilus* Membrane Vesicles

MVs from *L. acidophilus* MTCC 10307 were isolated using the ultracentrifugation procedure. Late log phase bacterial cultures were centrifuged at 5000 rpm for 20 minutes to remove cells. The step was repeated twice to remove extracellular debris and filtered through 0.45 and 0.22 µm filter and proceeded for ultracentrifugation at 1,50,000 rpm for 120 minutes. Then the pellet was resuspended in PBS and ultracentrifuged again at the same condition to obtain the MVs. The final MV preparation was stored at −80 °C until further use (13).

### Characterization of MVs using NTA

The *L. acidophilus* MVs were diluted in 0.22 µm filtered PBS and analyzed for particle size and concentration using a Malvern Panalytical NTA NS300 instrument. The instrument was operated under the following conditions: temperature, 25 °C; viscosity, 0.9 cP; dilution factor, 1:25; and syringe pump speed, 25 (14).

### Cell line source, maintenance, and chemicals

The RAW 264.7 macrophage cell line was obtained from the National Centre for Cell Sciences (NCCS), Pune, India. The cells were routinely maintained at 37 °C in DMEM high-glucose medium supplemented with 10% FBS under a humidified atmosphere containing 5% CO₂. All chemicals were purchased from HiMedia, India, unless otherwise stated (15).

### Cell viability assay

Cytotoxicity screening was performed for *L. acidophilus* MVs. RAW 264.7 cells were grown to approximately 90% confluence under the conditions described above in a 96-well plate. The cells were treated with different concentrations of MVs (3.3 × 10⁹ to 1.03 × 10⁸ particles/mL) for 24 h. Following treatment, 100 µL of MTT solution (0.5 mg/mL) was added to each well and incubated for 4 h. The resulting blue coloured formazan crystals were dissolved in 100% DMSO, and absorbance was measured at 570 nm (16).

### Relative gene expression – qRT-PCR

The relative gene expression of key inflammatory markers such as *Il1β*, *Il6*, and *Inos* was analyzed using qRT-PCR. Cells were grown to 90% confluence in 6 well plates. Inflammation was induced using LPS (1 µg/mL final concentration), followed by treatment with *L. acidophilus* MVs at concentrations of 2.1 × 10⁸ and 4.13 × 10⁸ particles/mL along with proteinase K treated *L. acidophilus* MVs at 4.13 × 10⁸ particles/mL. Diclofenac (10 µM) was used as a positive control. After 24 h, total RNA was isolated using the Macherey-Nagel TriPep RNA isolation kit according to the manufacturer’s protocol. RNA was reverse transcribed into cDNA using the ABclonal cDNA synthesis kit. qRT-PCR was carried out using Takara TB Green Premix Ex Taq II in a Bio-Rad CFX96 real-time PCR system. GAPDH was used as the reference gene, and relative gene expression was calculated using the 2^⁻ΔΔCt^ method (17).

### Nitric Oxide production assay

NO production was quantified using the Griess Reagent System (Promega, USA) by measuring nitrite (NO₂⁻) levels according to the manufacturer’s instructions. Cells were stimulated with LPS and treated with *L. acidophilus* MVs at concentrations of 2.1 × 10⁸ and 4.13 × 10⁸ particles/mL along with proteinase K treated *L. acidophilus* MVs at 4.13 × 10⁸ particles/mL. Diclofenac (10 µM) served as the positive control. After 24 h of treatment, culture supernatants were collected and centrifuged at 10,000 rpm for 5 min at 4 °C to remove extracellular debris. For nitrite estimation, 50 µL of clarified supernatant was mixed with 50 µL of sulfanilamide solution in a 96-well plate and incubated for 10 min at room temperature. Afterward, 50 µL of N-(1-naphthyl) ethylenediamine dihydrochloride (NED) solution was added and allowed to incubate for an added 10 min in the absence of light. Absorbance was measured at 540 nm using a microplate reader. Nitrite concentration was calculated from a sodium nitrite standard curve (0–100 µM) and expressed as µM nitrite (18).

### Reactive Oxygen Species production assay

Intracellular ROS generation was assessed using the fluorescent probe 2′,7′-dichlorodihydrofluorescein diacetate (DCFDA) followed by flow cytometric analysis. Cells were stimulated with LPS and treated with *L. acidophilus* MVs at concentrations of 2.1 × 10⁸ and 4.13 × 10⁸ particles/mL. Diclofenac (10 µM) was used as the positive control. After treatment, cells were gently washed with PBS and incubated with DCFDA (10 µM) for 30 min at 37 °C in the dark. Excess dye was removed by washing with PBS, and cells were resuspended in PBS. Fluorescence was immediately measured using a flow cytometer, with DCFDA detected in the FITC channel. A minimum of 10,000 events was acquired per sample (19).

### Western blot analysis

Western blot analysis was performed for IL1β, NFκB and its activated form NFκB p65 to correlate protein expression with the pro inflammatory cytokine profiles obtained from qRT-PCR. Cells were treated with RIPA buffer supplemented with protease inhibitor for 15 mins at 4°C. The lysate was centrifuged at 12000 rpm for 20 mins at 4°C to remove the debris and the lysate was mixed with Laemmli buffer, containing 2% sodium dodecyl sulfate, 10% glycerol, 2% β-mercaptoethanol, 50 mM Tris-HCl, 12.5 mM ethylenediaminetetraacetic acid (EDTA), and 0.02% bromophenol blue, followed by heat treatment at 90°C for 10 minutes. The equal volume of denatured proteins from each treatment groups were separated on a Tris-glycine gel with a linear concentration gradient of 5-12%. Following electrophoresis, proteins were transferred onto a Bio-Rad nitrocellulose membrane using wet transfer for 105 mins at 100 V. The membranes were then blocked with 5% BSA and washed prior to incubation with primary antibodies against β-actin, IL1β, NFκB, and NFκB p65 (Abclonal). After washing, the membranes were incubated with anti rabbit secondary antibody (Abclonal) for 1 hour at room temperature and developed using ECL solution (Abclonal) for 2 minutes. Imaging was performed using the Vilber system with automatic exposure settings (20).

### Repeated-dose *in vivo* oral toxicity study

A repeated dose 14 day oral toxicity study with 28 day recovery was conducted in accordance with OECD guideline 407 with slight modifications. Male animals were divided into vehicle control, 20 mg/kg MV, and 500 mg/kg MV groups and treated with the respective doses of MVs for 14 days. Animals were sacrificed at the end of 14 days and the corresponding satellite recovery groups were sacrificed after 28 days. Body weight was recorded once weekly and plotted as a graph (21).

### Histopathology of vital organs using Hematoxylin & Eosin staining

Tissues from vital organs including the heart, liver, spleen, and kidney were fixed in 10% neutral buffered formalin for 48 h at room temperature. Samples were dehydrated through a graded ethanol series (70%, 95%, and 100%) for 5 min each, cleared in xylene, and embedded in paraffin wax. Paraffin blocks were sectioned at 4-5 µm thickness using a rotary microtome and mounted on glass slides. For H&E staining, sections were deparaffinized in xylene, rehydrated through descending ethanol concentrations, and stained with hematoxylin for 5 min, followed by eosin counterstaining for 1-2 min. Sections were then dehydrated, cleared, mounted with DPX, and examined under a light microscope (Olympus CX23, Japan) (22).

### Serum analysis

Serum biochemical parameters including alkaline phosphatase (ALP), alanine aminotransferase (ALT), aspartate aminotransferase (AST), total cholesterol, high-density lipoprotein (HDL), urea, creatinine, glucose, and total bilirubin were analyzed using Arkray Autospan kits according to the manufacturer’s instructions to evaluate potential toxicological effects (23).

### Paw edema inflammatory model

The anti-inflammatory potential of *L. acidophilus* MVs was evaluated using an LPS induced acute paw edema model in Wistar albino rats. Paw edema was induced by intraplantar injection of 50 µL of LPS (50 µg) per animal. Subsequently, 6.7 × 10⁸ MVs per animal were administered, along with diclofenac (10 mg/kg) as the positive control. Paw thickness was measured using a vernier calliper at 0, 2, 4, 12, and 24 h (24). All animal experiments were approved by the Institutional Animal Ethics Committee (IAEC) and conducted in accordance with CPCSEA guidelines, Government of India (Approval number: VIT/IAEC/24/July23/05).

### Relative gene expression in paw tissue

Relative gene expression of *il1β*, *il6*, and *inos* from paw tissue was analyzed as described earlier. Paw tissue was homogenized using a mortar and pestle under liquid nitrogen prior to RNA isolation (25).

### Histopathology of paw tissue

Histopathological analysis of paw tissue was performed as described earlier using H&E staining to evaluate inflammatory changes between control and treatment groups (22).

### Proteomics analysis

Proteomic analysis of *L. acidophilus* MVs was performed using Thermo Fisher LC-MS/MS. Identified proteins were mapped against the *L. acidophilus* (26).

## Statistical analysis

All data was expressed as mean ± SD of three biologically independent experiments. Statistical significance was determined using one-way ANOVA followed by Tukey’s post hoc test, with p < 0.05 considered significant using GraphPad Prism 8.

## Results

### *In vitro* Anti-inflammatory Activity

#### Cell Culture and Cytotoxicity Assay

RAW 264.7 murine macrophage cells were maintained in DMEM high glucose medium supplemented with 10% FBS at 37 °C in a humidified atmosphere containing 5% CO₂. Cytotoxicity of *L. acidophilus* MVs was evaluated using MTT assay at concentrations ranging from 1.65 × 10⁹ to 1.03 × 10⁸ particles/mL **(Figure 1A-B)**. No significant reduction in cell viability was observed across the tested concentrations **(Figure 1B)**. Based on these results, 4.13 × 10⁸ and 2.1 × 10⁸ particles/mL were selected for subsequent experiments.

**Figure 1.**
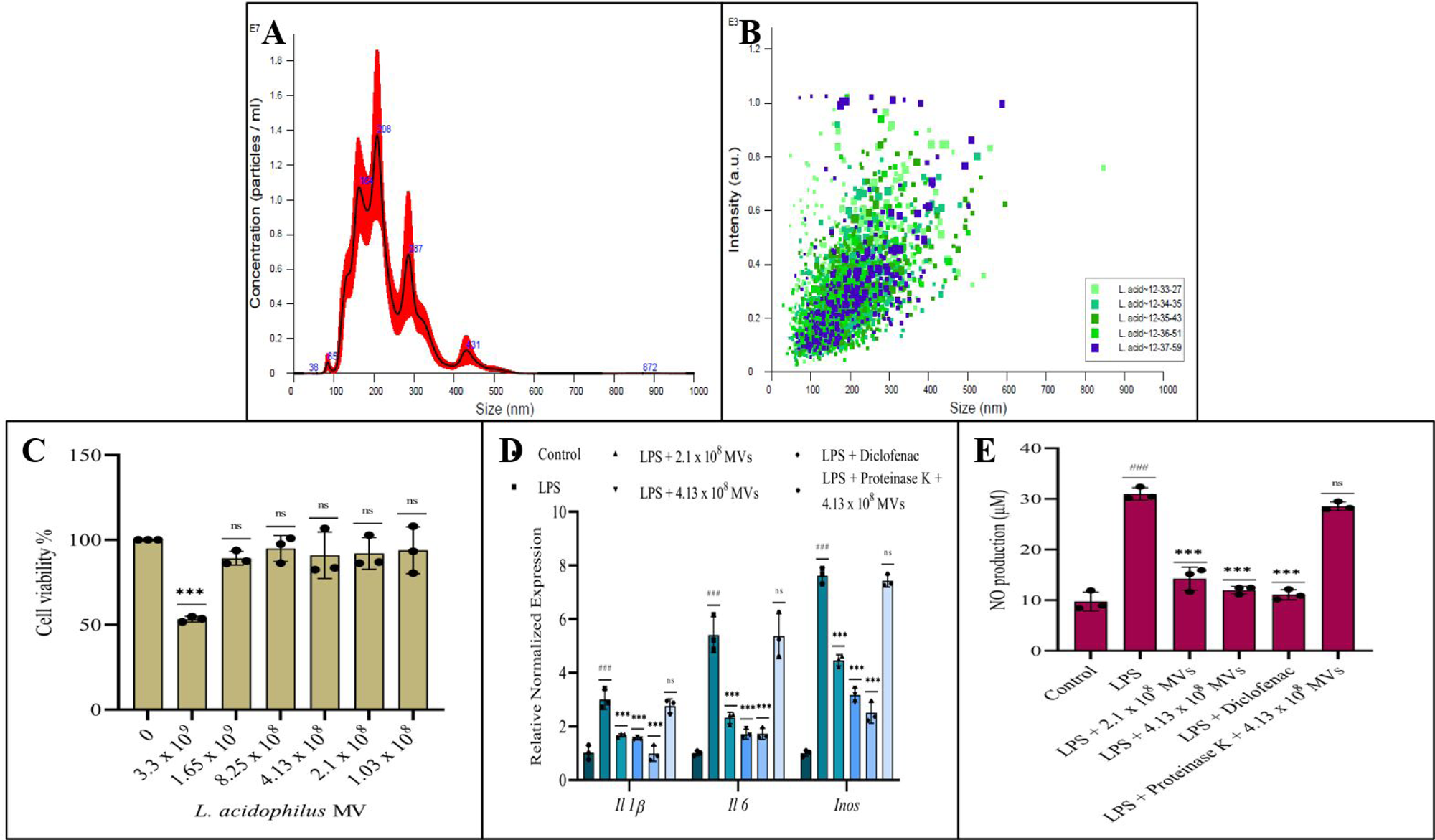
A) Concentration analysis in NTA revealed MV concentration to be 1.64 x 10^9^ particles/mL B) Dot plot representing the size distribution of the MVs in NTA C) Cell viability assay for *L. acidophilus* MVs at the concentration range of 3.3 x 10^9^ to 1.03 x 10^8^ particles/mL against RAW 264.7 macrophages D) Relative gene expression analysis of *Il 1β*, *Il 6* and *Inos* in control, LPS and MV treatments, reveals downregulation of the cytokines in MV treatment but reversed in the proteinase K treated MVs E) NO production assay revealed the downregulation in MV treatment at the concentrations of 2.1 and 4.13 x 10^8^ particles/mL. (***p < 0.001, ns p > 0.05)

#### Effect of MVs on Pro-inflammatory Gene Expression

Cells were stimulated with LPS (1 µg/mL final concentration) and treated with *L. acidophilus* MVs or diclofenac (10 µM) as a positive control. RT-qPCR analysis showed significant upregulation of *Il-1β*, *Il-6*, and *iNOS* following LPS induction. Treatment with *L. acidophilus* MVs significantly attenuated the expression of these genes in a concentration dependent manner. *Il-1β* expression was reduced by 1.8 and 2 folds at low and high MV concentrations, respectively. *Il-6* expression was reduced by 2.33 and 3.16 folds, while *iNOS* expression decreased by 1.7 and 2.4 folds compared to the LPS control. Interestingly proteinase K treated MVs did not reverse the upregulated cytokine expression **(Figure 1C)**.

#### Nitric Oxide Production

Nitric oxide production was assessed using the Griess assay. Consistent with reduced *Inos* expression as evaluated in qRT-PCR, MV treated groups portrayed significantly lower NO levels compared to the LPS-induced control and were comparable to the uninduced control. The percentage inhibition in the low, high and positive control diclofenac treated groups were 54, 61.29 and 64.19 respectively. Here, the proteinase K treated MV didn’t have the effect as the untreated one **(Figure 1D)**.

#### ROS Production Analysis

Intracellular ROS levels were measured using DCFDA staining followed by flow cytometry. LPS stimulation increased ROS levels to 88.22%, whereas MV treatment reduced ROS production to 48% and 40% at lower and higher concentrations, respectively. Diclofenac treatment reduced ROS levels to 32%. The uninduced control exhibited baseline ROS levels of 20.17% **(Figure 2A-F)**.

**Figure 2.**
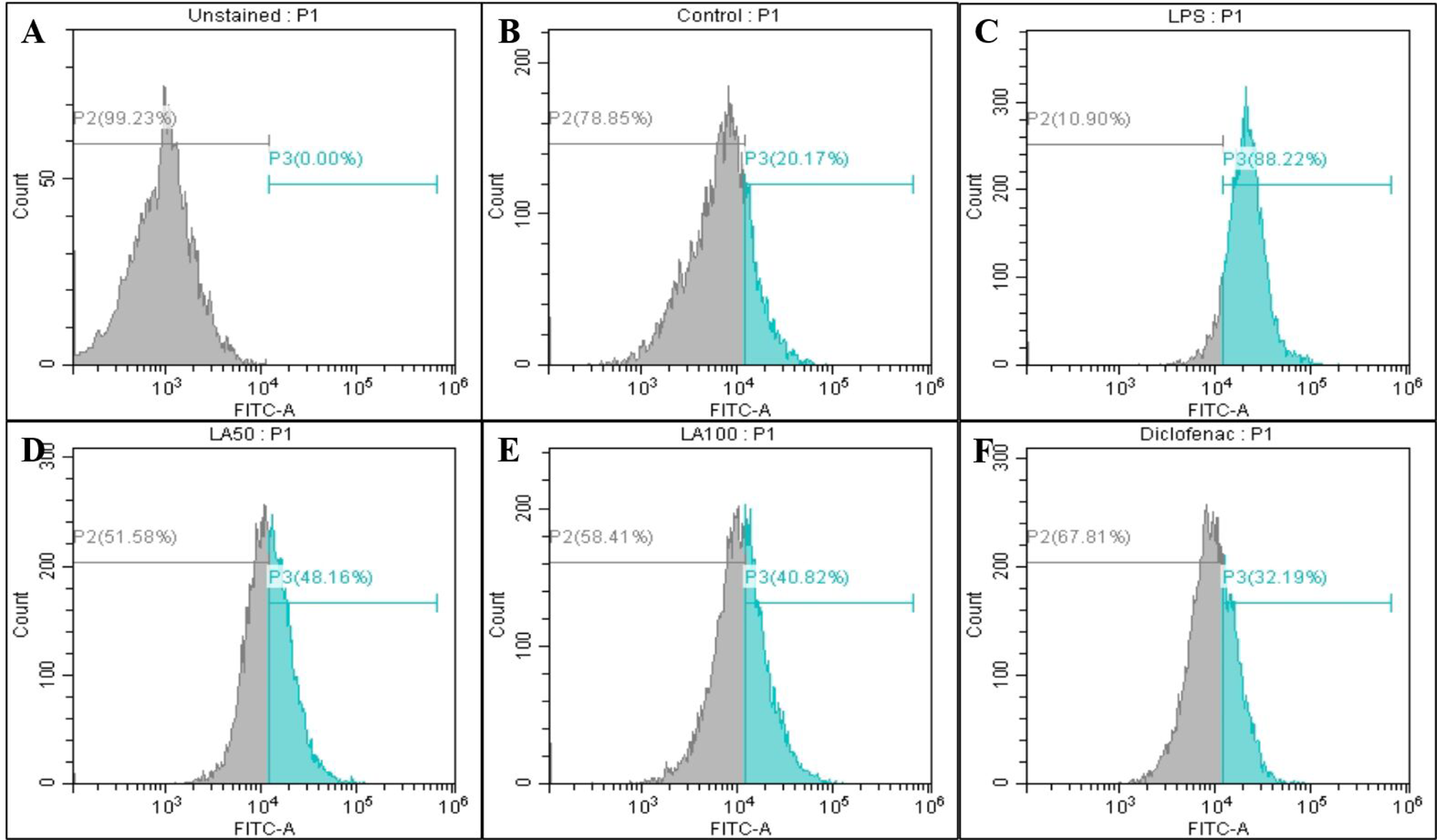
ROS production analyzed in flow cytometry with percentage of cells positive for ROS production A) Unstained cells B) control with 20.17% C) LPS with 88.22% D) *L. acidophilus* MVs at 2.1 x 10^8^ particles/mL with 48.16%, E) *L. acidophilus* MVs at 4.13 x 10^8^ particles/mL with 40.82% and F) positive control diclofenac with 32.19%.

#### Western blot analysis

The western blot analysis revealed the expression of NFκb-p65 and IL1β expression were upregulated in the LPS group which was reversed in the MVs treatment group. The expression of IL1β, NFκb-p65 and NFκb were normalized with the β-actin. The expression of NFκb-p65 was again compared with the normalized NFκb **(Figure 3A-C)**.

**Figure 3.**
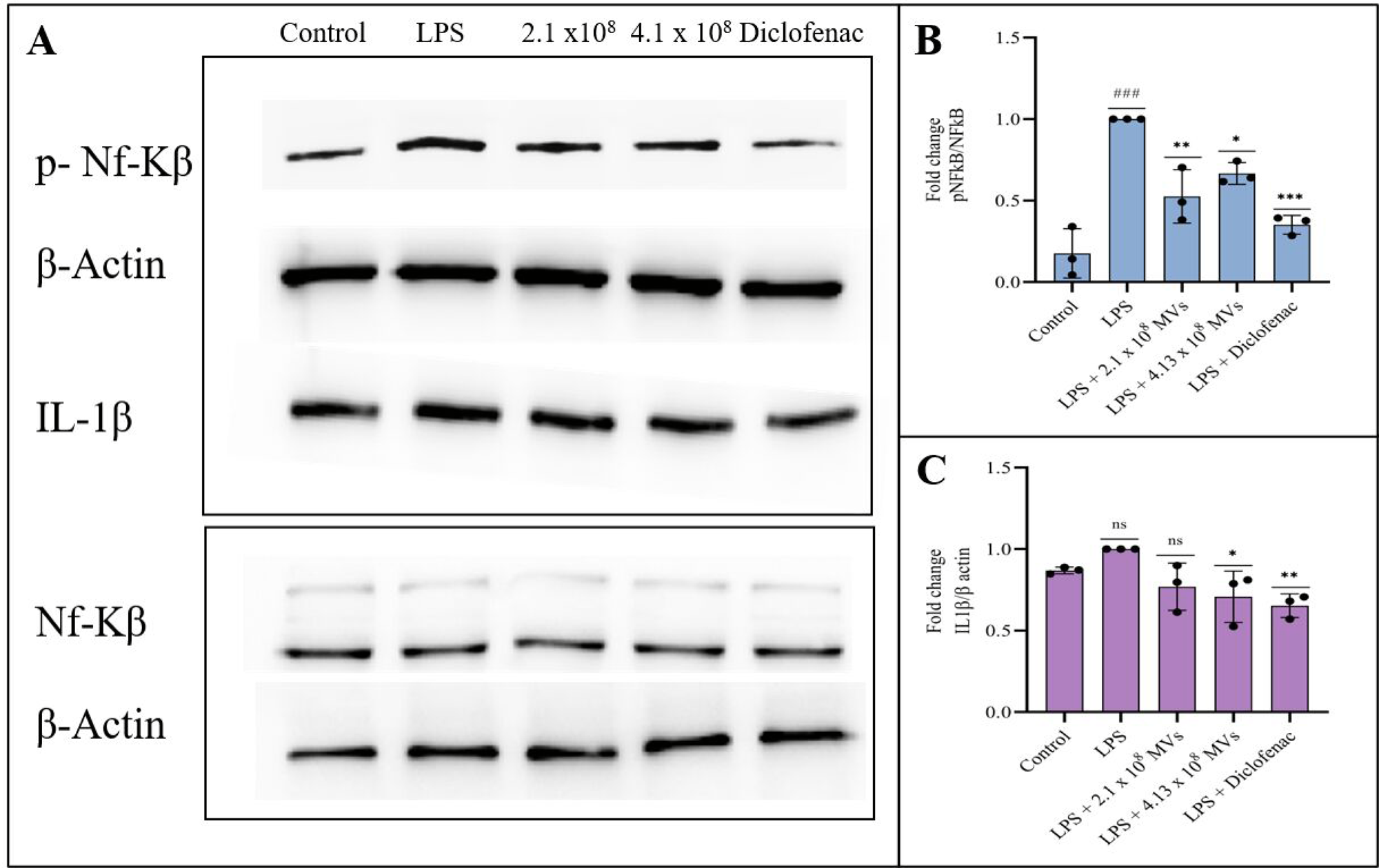
A) Western blot bands for proteins NF-kβ, p-NF-kβ & IL-1β B) Normalized value of p-NFkB against NFkB revealed reversal in the MV treatment group when compared to the LPS induction C) Normalized value of IL1β against β actin revealed reduction in the MV treatment when compared to the LPS. (***p < 0.001, **p < 0.01, *p < 0.05 ns p > 0.05)

### *In Vivo* Toxicity Evaluation

#### Repeated-Dose Oral Toxicity Study

A repeated dose 14 day oral toxicity study with 28 day recovery was conducted in accordance with OECD guideline 407. Animals were divided into vehicle control, 20 mg/kg MV, and 500 mg/kg MV groups, along with corresponding satellite recovery groups. No mortality or treatment related clinical signs were observed.

#### Body Weight, Serum Biochemistry, and Histopathology

Body weight progression remained comparable across all groups **(Figure 4)**. Serum biochemical parameters were estimated to assess any associated systemic toxicity following repeated oral administration of MVs derived from *L. acidophilus* in Wistar Albino rats. Across all experimental groups, including vehicle control, 20 mg/kg MV, 500 mg/kg MV, and satellite recovery groups, serum parameters showed only minimal and negligible variations. No dose dependent or treatment related alterations were observed. All measured serum parameters endured within normal physiological ranges, indicating the absence of MV induced hepatic, renal, or metabolic dysfunction. Furthermore, serum biochemical profiles of satellite recovery groups were comparable to those of the respective control groups, suggesting no delayed or reversible toxic effects **(Figure 5A-F)**. Histopathological examination was performed on heart, liver, spleen, and kidney tissues collected from control, MV treated, and satellite recovery groups. Heart sections from MV treated animals showed normal cardiac architecture with intact cardiomyocytes. Clearly defined myonuclei were observed, and intercalated disc spaces appeared normal. No evidence of myocardial degeneration, necrosis, inflammatory cell infiltration, or vascular abnormalities was detected across treatment or recovery groups **(Figure 6A-J)**. Histological sections of liver tissue exhibited normal hepatic architecture in all MV treated and satellite groups. Hepatocytes appeared polygonal and well organized around a centrally located vein. Hepatic sinusoids were intact, and portal triad structures, including portal vein and bile ducts were normal. No signs of hepatocellular degeneration, inflammation, or fibrosis were observed **(Figure 7A-J)**. Spleen sections from MV treated animals showed well preserved splenic architecture. Distinct white pulp regions with centrally located arterioles and normal red pulp containing intact sinusoids were observed. There were no indications of lymphoid depletion, congestion, or abnormal cellular proliferation in any treatment group **(Figure 8A-J)**. Kidney sections revealed normal renal histology in MV treated and satellite groups. Glomeruli appeared intact with well-defined Bowman’s capsules, and renal tubules showed normal morphology. No evidence of tubular degeneration, glomerular congestion, interstitial inflammation or fibrosis was observed **(Figure 9A-J)**.

**Figure 4.**
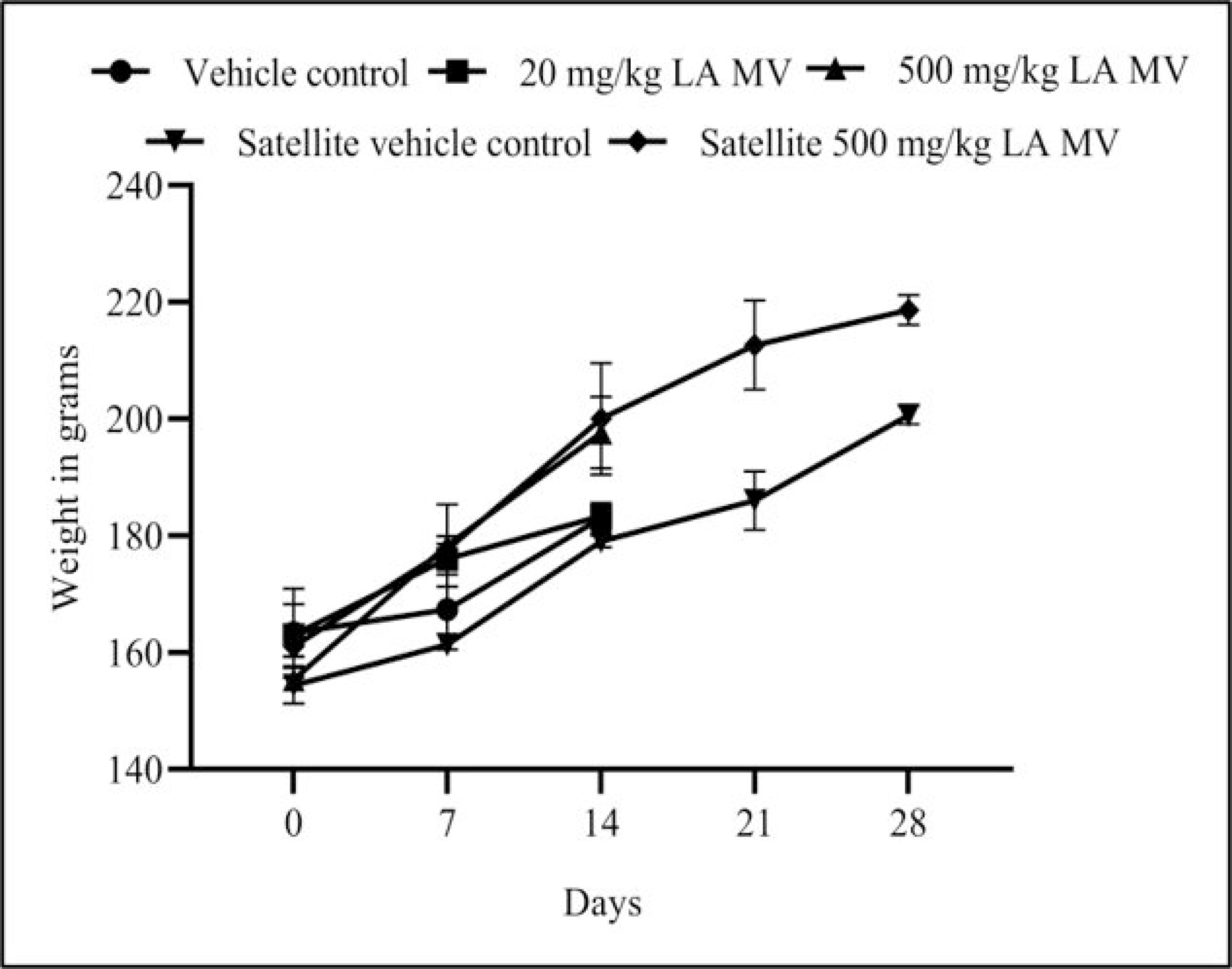
Bodyweight of rats from *in vivo* toxicity assay over a period of 28 days.

**Figure 5.**
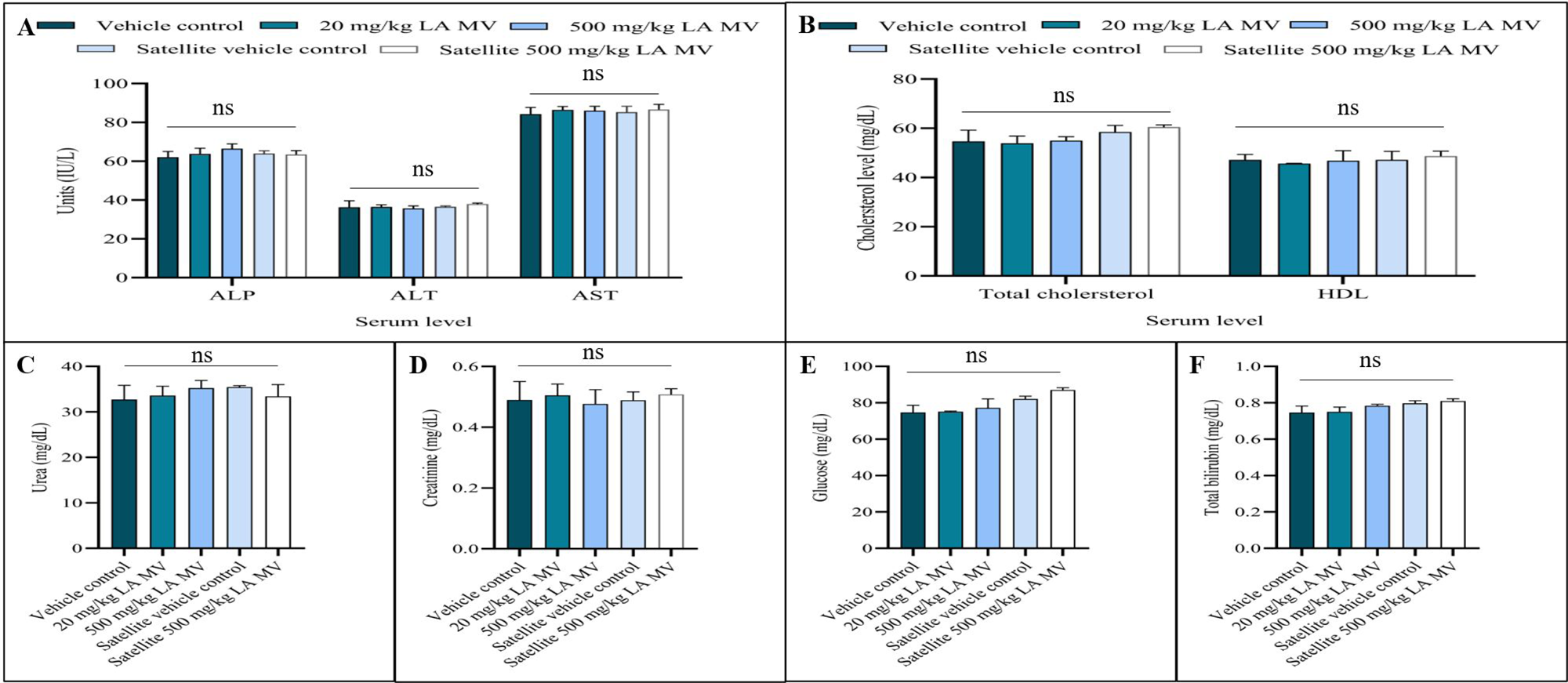
Serum analysis for toxicity studies A) ALP ALT and AST B) Total cholesterol and LDL C) Urea D) Creatinine E) Glucose F) Total bilirubin. (ns p > 0.05)

**Figure 6.**
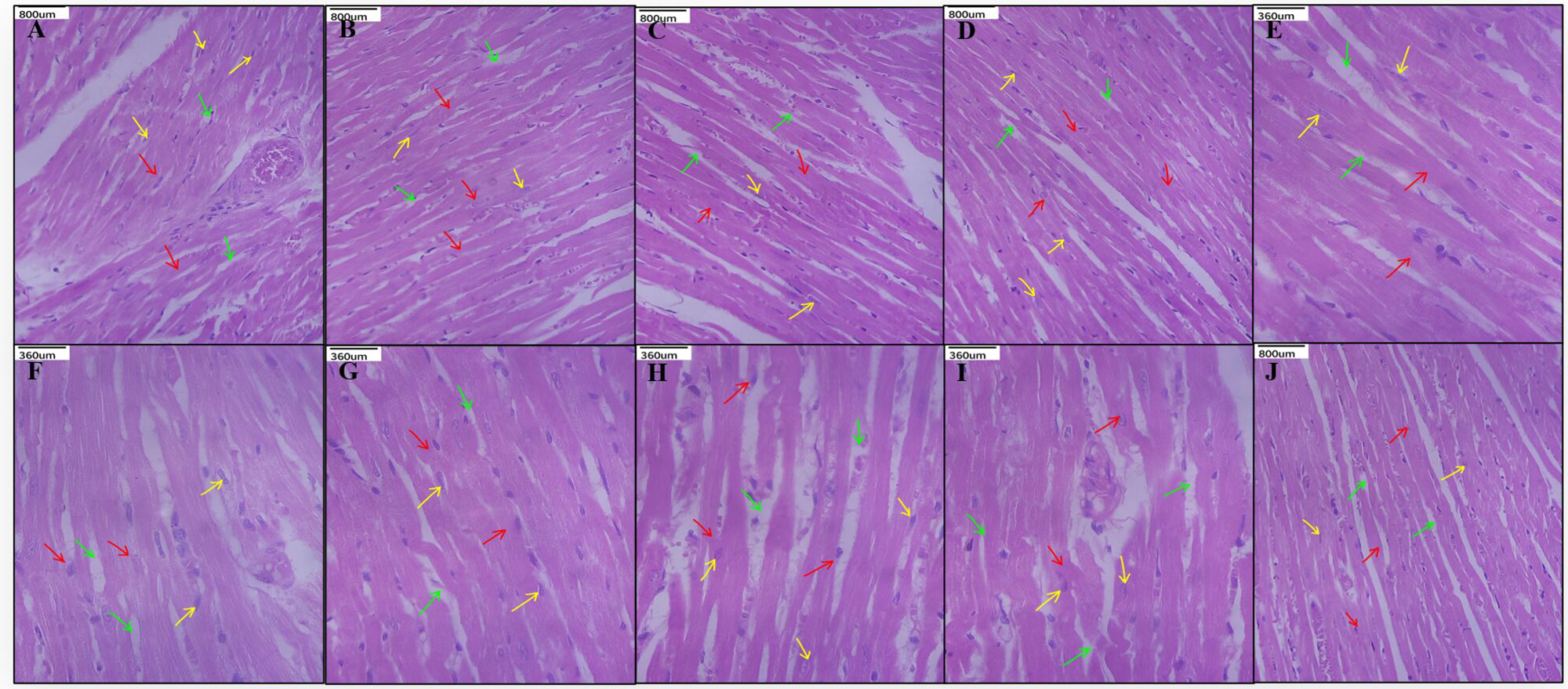
Histopathology analysis of toxicity studies, Heart A) Vehicle control B) 20 mg/kg MV C) 500 mg/kg MV D) satellite vehicle control E) Satellite 500 mg/kg MV.

**Figure 7.**
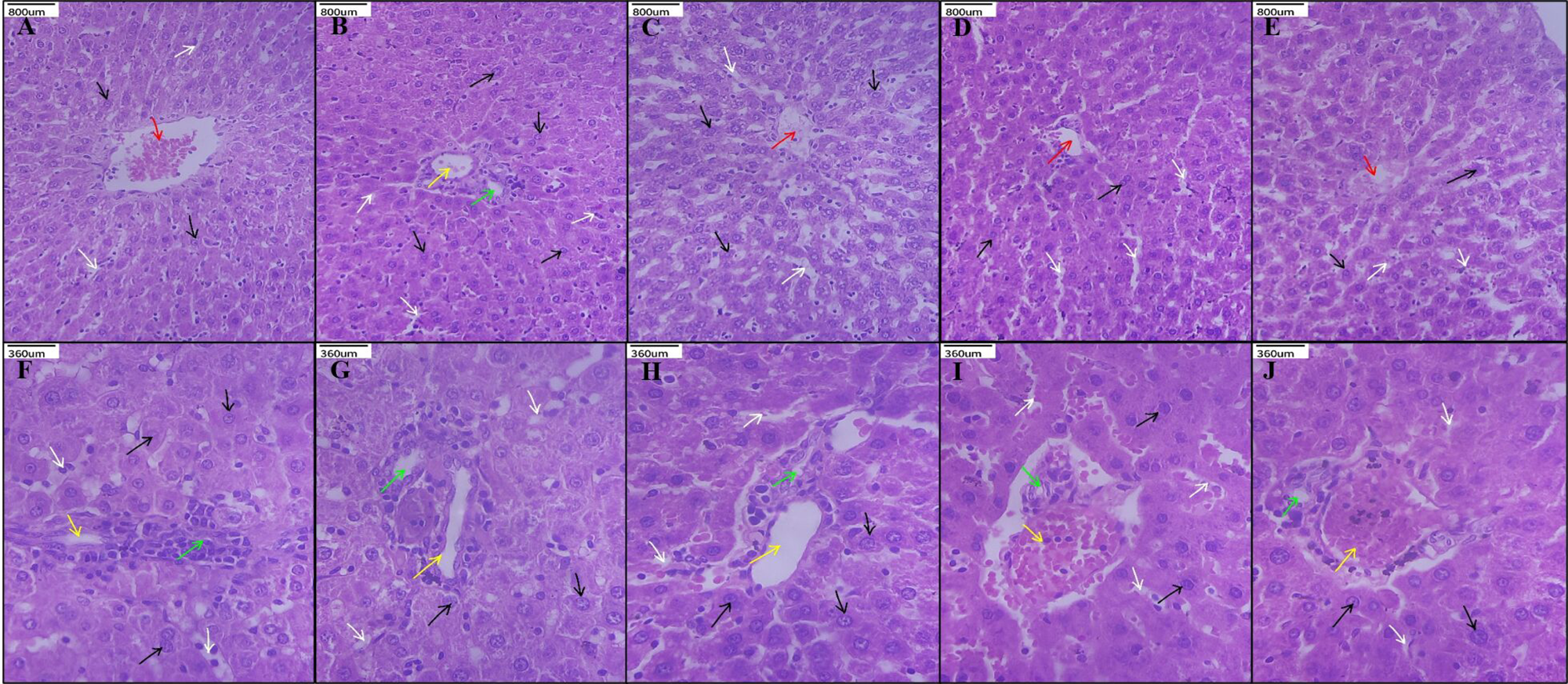
Histopathology analysis of toxicity studies, Liver A) Vehicle control B) 20 mg/kg MV C) 500 mg/kg MV D) satellite vehicle control E) Satellite 500 mg/kg MV.

**Figure 8.**
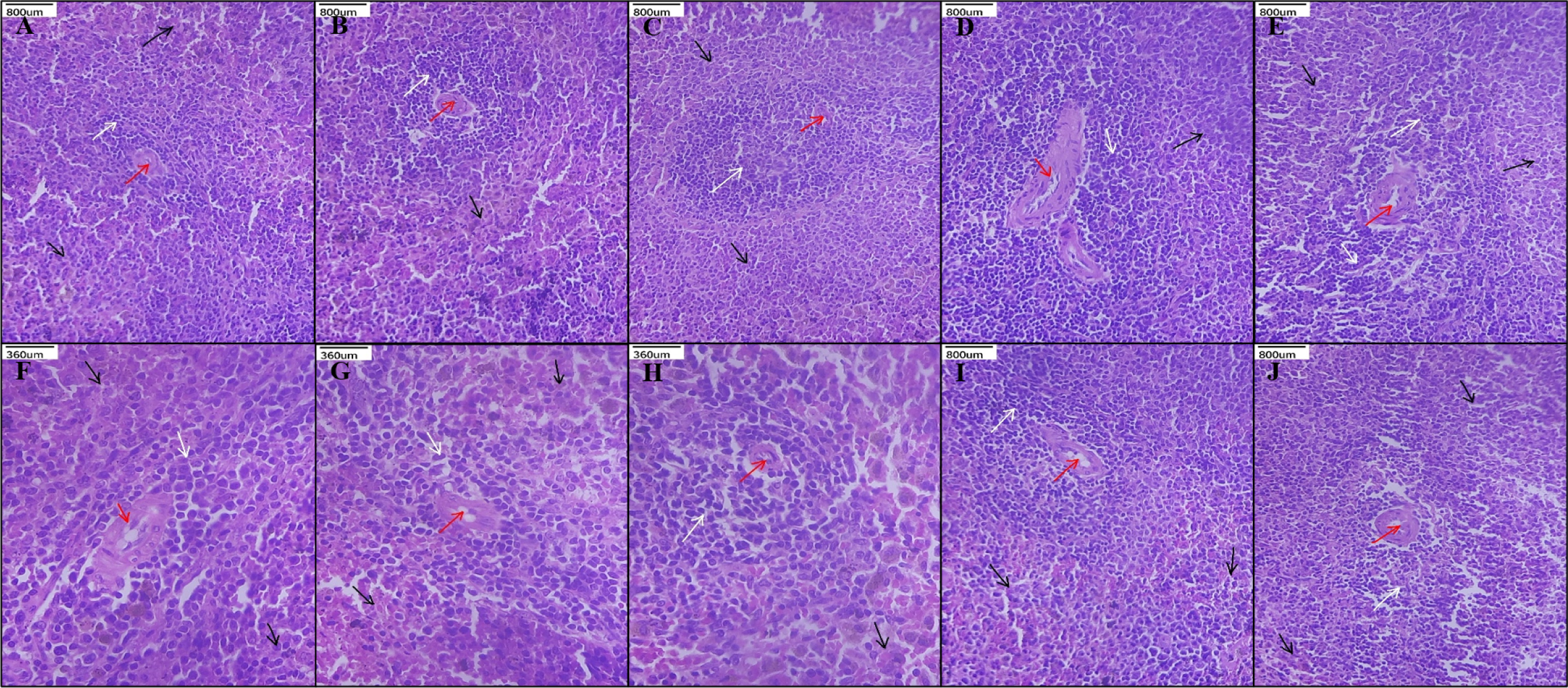
Histopathology analysis of toxicity studies, Spleen A) Vehicle control B) 20 mg/kg MV C) 500 mg/kg MV D) satellite vehicle control E) Satellite 500 mg/kg MV.

**Figure 9.**
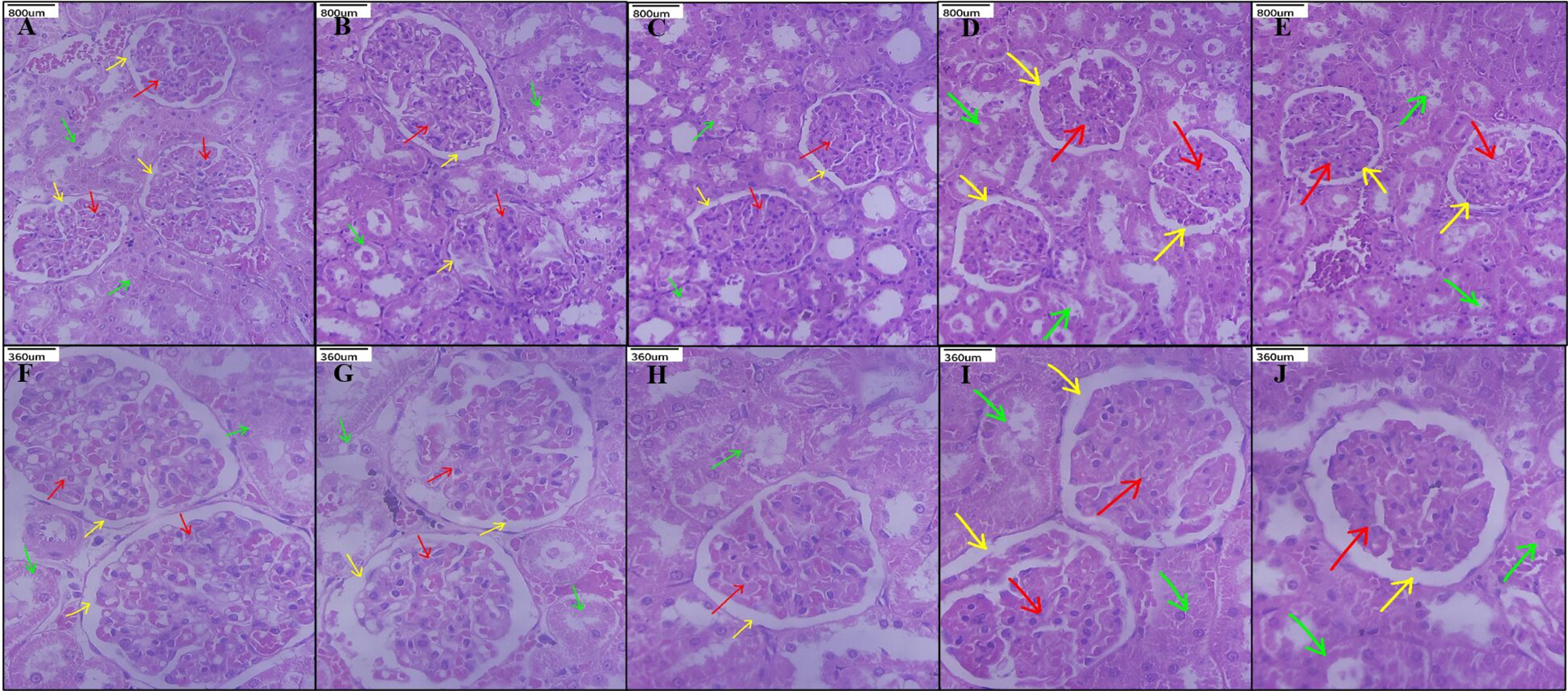
Histopathology analysis of toxicity studies, Kidney A) Vehicle control B) 20 mg/kg MV C) 500 mg/kg MV D) satellite vehicle control E) Satellite 500 mg/kg MV.

### *In Vivo* Anti-inflammatory Activity

#### Hind Paw Edema Model

Anti-inflammatory efficacy of MVs derived from *L. acidophilus* was evaluated using an LPS induced hind paw edema model in Wistar albino rats. LPS administration resulted in a marked increase in paw width in the untreated group, confirming successful induction of acute inflammation. In contrast, animals treated with *L. acidophilus* MVs exhibited a noticeable reduction in paw edema over the 24 hour observation period. The extent of edema resolution in the MV treated group was comparable to that observed in the diclofenac treated positive control group **(Figure 10A)**. No abnormal behavioural changes were noted during the experimental period. These observations indicate that MV treatment effectively reduced LPS-induced paw swelling.

**Figure 10.**
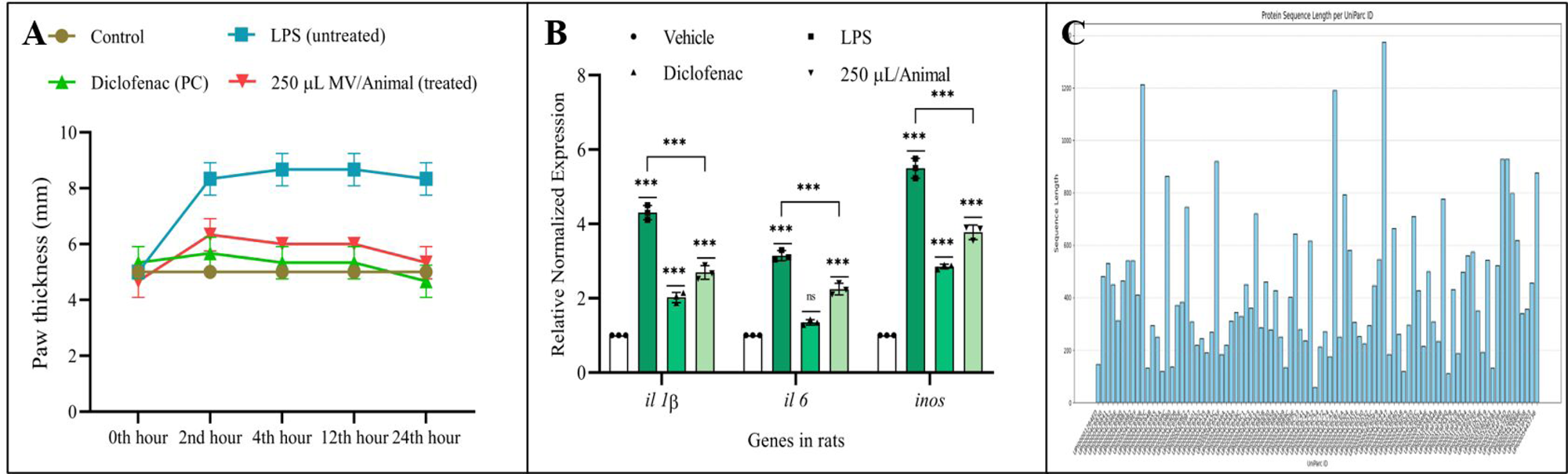
A) Paw width of animals shows reducing in the MV treatment group B) Relative expression of pro inflammatory cytokines shows downregulation of *Il1β*, *Il6* and *Inos* expression in the MV treatment group C) S-layer protein identification in MVs through LC-MS/MS. (***p < 0.001, ns p > 0.05)

#### Pro inflammatory Cytokine Expression in Paw Tissue

To assess inflammatory mediator expression at the molecular level, RT-qPCR analysis was performed on paw tissue collected from experimental animals. LPS treated animals showed marked upregulation of *il1β*, *il6*, and *inos* transcripts compared to the untreated control group. In contrast, paw tissues from MV treated animals demonstrated significant downregulation of these pro inflammatory genes when compared with the LPS group. A similar reduction in cytokine expression was observed in the diclofenac treated group. The inhibition percentages observed for *il1β*, *il6*, and *inos* in the MV treated group were consistent with the anti-inflammatory trend observed in the paw edema measurements **(Figure 10B)**.

#### Histopathological Analysis of Paw Tissue

Histopathological evaluation of paw tissue sections was performed using hematoxylin and eosin staining to examine inflammation associated structural changes. Sections from the LPS treated group exhibited pronounced pathological alterations, including epidermal hyperkeratosis and hyperplasia, along with dense infiltration of inflammatory cells within the dermis and the presence of granulation tissue. In contrast, paw tissue sections from MV treated animals showed near normal histological architecture, characterized by an intact epidermis, organized dermal collagen bundles, and minimal inflammatory cell infiltration. Similar histological features were observed in the diclofenac treated group. These findings demonstrate that MV treatment effectively attenuated LPS induced inflammatory tissue damage **(Figure 11A-H)**.

**Figure 11.**
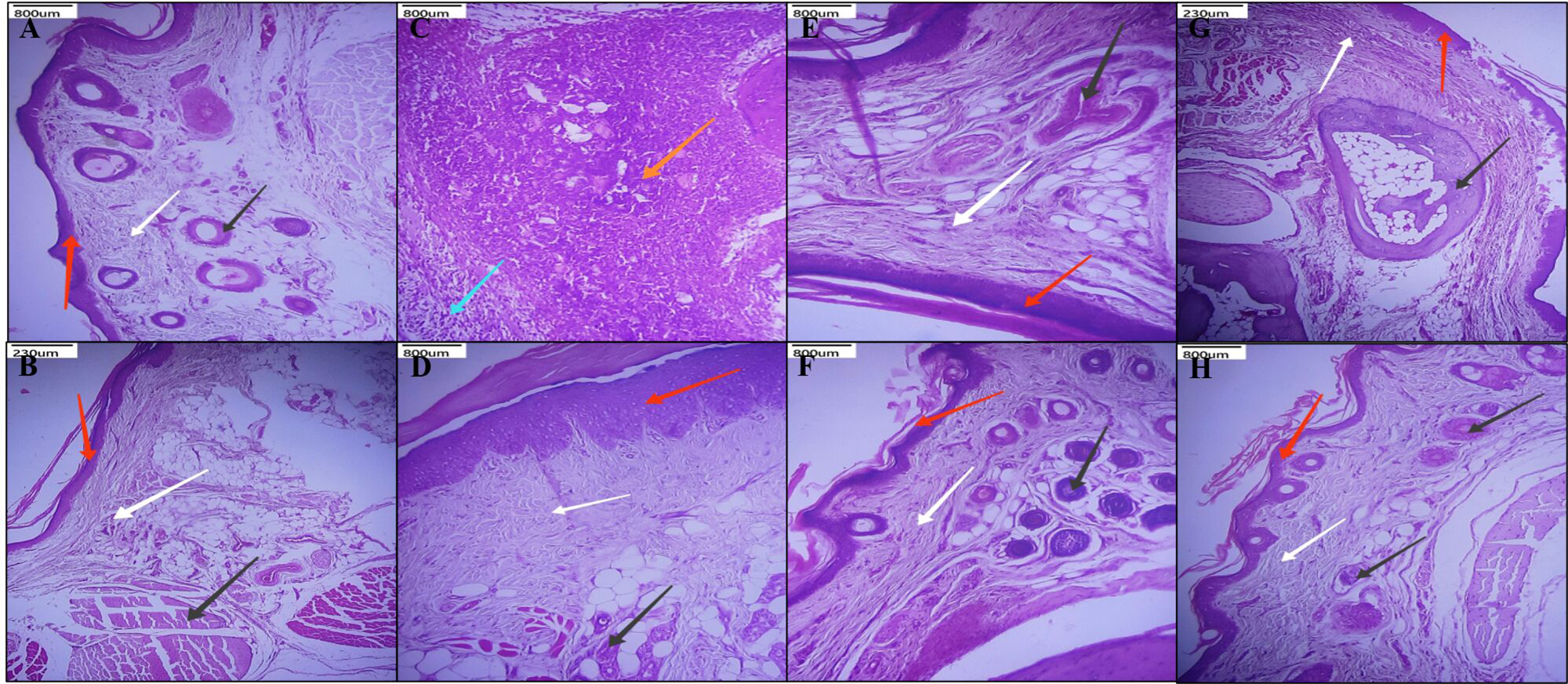
Histopathology analysis of paw edema in A-B) Control C-D) LPS E-F) 6.7 x 10^8^ MVs G-H) Diclofenac treatments groups show significant reduction in the paw width in the treatment groups when compared to the LPS.

#### Proteomic Identification of Bioactive Components

LC-MS/MS based proteomic analysis identified surface-layer protein A (SlpA) as a prominent component of *L. acidophilus* MVs, supporting its role in mediating anti-inflammatory effects via NFκB modulation **(Figure 10C)**.

## Discussion

The present study demonstrates that MVs derived from *L. acidophilus* MTCC 10307 exert potent anti-inflammatory effects while maintaining an excellent safety profile. By integrating *in vitro* macrophage based assays, repeated dose oral toxicity evaluation, and *in vivo* anti-inflammatory validation, this work suggests that *L. acidophilus* MVs as biologically active yet nontoxic immunomodulatory agents. Macrophages play a central role in orchestrating inflammatory responses through the production of cytokines and inflammatory mediators upon stimulation with bacterial endotoxins such as LPS (27). In this study, *L. acidophilus* MVs significantly attenuated LPS-induced upregulation of *Il1β*, *Il6*, and *Inos* in RAW 264.7 macrophages. These mediators are hallmark indicators of inflammatory activation and are directly involved in amplifying immune responses and tissue damage (28). Anti-inflammatory effects of *L. acidophilus* have been widely reported using whole cells, often administered in heat inactivated form, as well as cell free supernatants (29). While heat inactivation improves safety by eliminating bacterial viability, it still exposes the host immune system to the full structural complexity of the bacterial cell, including cell wall components capable of broad innate immune activation (30). In contrast, MVs represent a refined postbiotic platform, delivering a defined nano-sized and encapsulated cargo of bioactive molecules without the bulk cellular architecture or pH dependent metabolic byproducts associated with supernatants (31). This selective and controlled mode of delivery enables effective immunomodulation while minimizing nonspecific immune stimulation, thereby conferring an improved safety profile and enhanced translational potential for anti-inflammatory applications.

The observed reduction in *Inos* expression was functionally corroborated by a decrease in nitric oxide production, confirming that MV mediated transcriptional suppression translated into reduced inflammatory effector output. Nitric oxide, although beneficial at physiological levels, contributes to oxidative and nitrosative stress when excessively produced during inflammation (32). Therefore, the ability of *L. acidophilus* MVs to downregulate both *Inos* expression and NO production highlights their role in mitigating inflammation associated cellular stress. ROS act as secondary messengers in inflammatory signalling cascades and are strongly associated with macrophage activation and tissue injury (33). LPS stimulation resulted in a marked elevation of intracellular ROS levels, whereas MV treatment significantly reduced ROS accumulation in a concentration dependent manner. This reduction suggests that *L. acidophilus* MVs not only suppress inflammatory gene expression but also alleviate oxidative stress, which is closely linked to chronic inflammatory pathology. Proteinase K treated MVs mitigated the effect which explains that the proteins present in the MVs is responsible for the observed actions (8). The concurrent reduction of ROS and nitric oxide further indicates that MV treatment restores redox homeostasis under inflammatory conditions, thereby limiting cellular damage and sustaining macrophage viability. NF-κB is a master transcriptional regulator controlling the expression of numerous pro-inflammatory genes, including IL-1β, IL-6, and iNOS. The concordance between gene expression and functional assays are consistent with suppression of NFκB mediated signalling pathways (5).

A critical requirement for translational application of biologically derived therapeutics is a robust safety profile. The repeated dose oral toxicity study conducted in accordance with OECD guideline 407 demonstrated that *L. acidophilus* MVs did not induce any treatment related adverse effects, even at the highest tested dose. Body weight progression remained normal, serum biochemical parameters showed only negligible variations, and histopathological evaluation of major organs revealed intact tissue architecture. The inclusion of a recovery (satellite) group further confirmed the absence of delayed or reversible toxic effects. These findings collectively establish that *L. acidophilus* MVs are systemically well tolerated, supporting their suitability for prolonged or repeated administration. The anti-inflammatory efficacy of *L. acidophilus* MVs was further validated using an LPS induced hind paw edema model, a well-established experimental system for evaluating acute inflammation (24). MV-treated animals exhibited significant resolution of paw edema over time, comparable to the standard anti-inflammatory drug diclofenac (34). Consistent with *in vitro* observations, RT-qPCR analysis of paw tissue demonstrated reduced expression of *il1β*, *il6*, and *inos* in MV treated animals. Histopathological examination provided structural confirmation of inflammation resolution, showing normalized epidermal architecture and reduced inflammatory cell infiltration in MV treated groups. The strong correlation between molecular, morphological, and functional outcomes underscores the reproducibility and robustness of MV-mediated anti-inflammatory effects.

Proteomic analysis identified surface-layer protein A (SlpA) as a prominent component of *L. acidophilus* MVs. SlpA has been previously reported to exert anti-inflammatory effects by suppressing pro-inflammatory mediators such as iNOS, COX-2, TNF-α, and IL-1β in LPS-stimulated macrophages. It was previously demonstrated that SlpA modulates inflammation through regulation of TLR4-dependent MAPK and NF-κB pathways, as well as NOD2/NLRP3-associated signalling (35, 36).

The identification of SlpA within *L. acidophilus* MVs in the present study provides a plausible molecular basis for the observed suppression of NF-κB activation and downstream inflammatory mediators. While the present work does not isolate SlpA independently, its presence strongly suggests that MV associated SlpA contributes, at least in part, to the anti-inflammatory activity observed. However, additional studies employing SlpA depletion or recombinant SlpA would be required to definitively establish its causal role in MV mediated immunomodulation. Taken together, these findings demonstrate that *L. acidophilus* MVs exert a multi-layered anti-inflammatory effect by suppressing inflammatory signalling pathways, reducing oxidative and nitrosative stress, and resolving tissue inflammation without inducing systemic toxicity. The convergence of *in vitro* mechanistic data and *in vivo* functional outcomes highlights the therapeutic promise of probiotic derived MVs as safe, biologically active alternatives to conventional anti-inflammatory agents. The novelty of this study lies in demonstrating that MVs derived from *L. acidophilus* retain potent immunomodulatory activity while eliminating the safety concerns associated with live probiotic administration. By combining macrophage-based mechanistic assays, OECD-guided toxicity evaluation, and *in vivo* inflammation resolution, this work provides a translational framework for probiotic-derived vesicles as next-generation anti-inflammatory therapeutics. These findings advance probiotic based therapeutics beyond live bacterial formulations toward safer, well defined, vesicle based immunomodulatory interventions.

## Conclusion

MVs derived from *L. acidophilus* exhibit robust anti-inflammatory activity *in vitro* and *in vivo* while maintaining an excellent safety profile. These findings highlight their potential application as safe, probiotic derived immunomodulatory agents for the management of inflammatory conditions. This study is among the first to demonstrate that MVs from *L. acidophilus* exert robust anti-inflammatory effects *in vivo* while maintaining an excellent safety profile upon repeated oral administration. The identification of SlpA within these vesicles further provides mechanistic insight into their immunomodulatory action, positioning probiotic derived MVs as promising, cell-free alternatives to conventional anti-inflammatory agents.

## Acknowledgment

The authors also thank the Vellore Institute of Technology for the basic and advanced infrastructure facilities. The authors thank Ms. Huldah Pearlin Sarah Lazarus of the School of Biosciences and Technology, Vellore of Institute of Technology, for her valuable support during the execution of western blotting and animal experiments.

## Author contributions

M.V and E.N designed the study. M.V performed all the experiments and wrote the manuscript. E.N finalized and edited the manuscript. E.N supervised all the works performed.

## Data availability statement

The datasets utilized and/or analysed during the present study are available from the corresponding author upon reasonable request.

## Ethics approval statement

This study was carried out in compliance with the Animal Research: Reporting In Vivo Experiments (ARRIVE) guidelines and was granted ethical approval by the institutional animal ethical committee of Vellore Institute of Technology (Approval number: VIT/IAEC/24/July23/05)

